# STED/AFM as a tool to investigate mechanical and adhesive properties of migrating keratinocytes

**DOI:** 10.1101/2025.09.10.675301

**Authors:** Mariya Y. Radeva, Jens Waschke, Michael Fuchs

## Abstract

E-cadherin is a central junctional molecule of adherens junctions that regulates epithelial integrity. In keratinocytes, wound healing critically depends on dynamic modulation of adhesion and cytoskeletal organization. Here, we applied wound healing assay and the combination of the stimulated emission depletion (STED) microscopy with the atomic force microscopy (AFM) (STED/AFM hybrid technique), to study E-cadherin binding properties during keratinocyte migration. Wound closure occurred within 6 h, an effect associated with E-cadherin accumulation at the leading edge. For location-specific binding measurements with the STED/AFM technique, we overexpressed E-cadherin in mouse keratinocytes and performed single molecule force spectroscopy measurements. Inhibition of actin polymerization abolished E-cadherin binding and further reduced overall cellular stiffness. To study adhesion mechanics during migration, we used the STED/AFM technique and performed single-molecule force spectroscopy at the leading edge of keratinocytes migrating into cell-free areas, generated by removal of two-well inserts. This approach enabled, for the first time, simultaneous measurements of single-molecule binding properties and cellular mechanistic properties in actively migrating keratinocytes. Our results revealed that E-Cad molecules present during migration exhibit binding properties comparable to those of E-Cad in stable AJs. This suggests that these molecules remain functional competent and may be readily available for rapid re-engagement in cell-cell adhesion when required. We introduce a powerful methodology to investigate single molecule binding properties in migrating cells, offering new opportunities to analyze epithelial repair at molecular resolution.

## Introduction

E-Cadherin (E-Cad) is a transmembrane glycoprotein that mediates calcium-dependent intercellular adhesion and is a core component of the adherens junction (AJ) complex [1]. The primary function of AJs is to maintain intercellular adhesion and provide resistance to external mechanical forces. E-Cad connects to the actin cytoskeleton via intracellular plaque proteins [2, 3]. Cell-cell adhesion mediated by AJs is a highly dynamic process, playing a crucial role during epithelial-to-mesenchymal transition (EMT), both during normal development and along carcinogenesis [3, 4]. In adult tissues, the loss of E-Cad is a key event in epithelial tumorigenesis and is observed across a wide range of tumor types. Conversely, re-expression of E-Cad in cancer cells has been shown to prevent tumor progression [5, 6]. In cancer, reduced E-Cad expression is a hallmark of the invasive cell phenotype [7].

Skin wound healing is a highly intricate process that typically progresses through three overlapping phases: inflammation, tissue formation, and remodeling [8, 9]. During the tissue formation phase, in addition to angiogenesis and extracellular matrix deposition, keratinocytes migrate across the fibrin matrix initially settled at the base of the wound to re-establish the epidermal barrier, an essential step in wound closure both *in vitro* and *in vivo* [10]. At the wound edge, keratinocytes undergo a partial EMT, enabling them to regain migratory capacity. In this state, E-Cad expression is downregulated through activation of the epidermal growth factor (EGF) receptor, which in turn reduces intercellular adhesion and facilitates keratinocytes to migrate across the wound bed [11]. Interestingly, E-Cad is not completely lost during this process. Instead, the remaining E-Cad molecules sustain a degree of cell-cell adhesion, supporting a mechanism known as collective migration [12]. However, in migrating keratinocytes, the binding properties and functional role of the leftover E-Cad molecules remain poorly understood.

To investigate both the mechanical and the binding properties of E-Cad in migrating keratinocytes, we performed scratch wound assays, immunostaining analysis and a novel hybrid approach that combines STED microscopy with AFM single-molecule force spectroscopy (SMFS) [13, 14]. Using this STED/AFM system, we were able to measure specific E-Cad adhesion events on living, migrating keratinocytes.

Our results show that E-cadherin molecules present during migration retain binding properties comparable to those in stable adherens junctions, indicating they remain functional and available for rapid re-engagement in cell–cell adhesion.

## Materials and methods

### Cell culture, transfection and reagents

Wild-type murine keratinocytes (MEKs) were isolated and cultured as previously described [15]. Cells were maintained in complete FAD medium (ThermoFisher Scientific, Germany) and initially grown in low-calcium medium (0.05 mM CaCl₂). Upon reaching confluency, the culturing conditions were switched, and the cells were cultivated under high-calcium conditions (1.8 mM CaCl₂) to induce differentiation. All experiments were conducted 24 hours after the calcium switch was initiated. For E-Cad overexpression, murine keratinocytes were grown to 70% confluency and transiently transfected with the pSNAPf-ECad-N plasmid using Lipofectamine 3000, following the manufacturer’s instructions (Invitrogen, Carlsbad, USA) [16]. A day after transfection, the cells were exposed to high-calcium medium for another 24 h and then subjected to experiments. Inhibition of actin polymerization was achieved by 1 hour pharmacological treatment with 2.0 µg/ml Latrunculin B (LatB; Merck, Darmstadt, Germany), as previously published [17].

### Wound assays

MEK cells were seeded into 24-well plates, with or without coverslips. After reaching confluency, cells were switched to high-calcium medium (1.8 mM Ca²⁺). 24 hours later a scratch was introduced into the cell monolayer using a sterile P200 pipette tip. Cell migration was monitored over time using bright-field microscopy (Axio Vert A1; Carl Zeiss AG, Oberkochen, Germany) or immunostaining was performed at 0, 2, 4, 6, and 8 h post scratch. For STED/AFM experiments, a sterile culture-Insert 2 well (Ibidi, Gräfelfing, Germany) was applied on the coverslip and removed 6 hours before the test performance, to create a cell-free gap for wound healing assays.

### Immunostaining, labelling of actin filaments and cell nuclei

MEK cells were fixed by incubation with ice-cold absolute ethanol for 30 minutes, followed by a 3-minute treatment with ice-cold acetone, both on ice. For detection of E-cad, a primary antibody directed against the protein was used (Invitrogen, Frankfurt, Germany). Actin filaments were visualized using Alexa 488-phalloidin (Dianova, Hamburg, Germany) and cell nuclei were stained by DAPI (Roche, Mannheim, Germany). For confocal microscopy, a Leica SP5 confocal microscopy with a 63 x NA 1.4 PL APO objective controlled by LAS AF software (Leica, Mannheim, Germany) was used. For E-Cad intensity values we used the whole region of interest and measured the integrated density values.

### Sample preparation for STED/AFM measurements

Staining of differentiated MEK’s monolayers was performed according to the manufacturer’s instructions. Briefly, cells exposed to complete high-calcium medium were incubated for 1hwith SiR-Actin (1 µM; Spirochrome, Denver, USA). After staining, the cells were washed once more with medium and transferred into the BioCell holder (Bruker Nano, Berlin, Germany), pre-heated to 37 °C, within the integrated STED/AFM setup.

### Stimulated emission depletion microscopy (STED)

Imaging was performed using the Expert Line STED microscope system from Abberior (Abberior Instruments GmbH, Göttingen, Germany), equipped with a 100 × oil immersion objective. Image acquisition was carried out using the Imspector software (Abberior). Fluorescent dye SNAP-CELL-TMR STAR was excited at 554 nm and the Sir-actin at 652 nm, using pulsed diode lasers (PiL063X, Advanced Laser Diode Systems). STED depletion was achieved with a pulsed 775 nm laser operated at 10 to 30% power, with a gating delay of 800 ps. Fluorescence emission was detected using an avalanche photodiode detector within the spectral ranges 650–720 nm.

### Atomic force microscopy (AFM)

AFM measurements were performed exclusively within the STED/AFM setup using a NanoWizard 4 AFM system (Bruker Nano, Berlin, Germany). Cantilever functionalization followed previously established protocols [18]. The recombinant protein was applied at 0.15 mg/ml concentration and covalently linked to a flexible Si₃N₄ cantilever (MLCT probes, nominal spring constant 0.03 N/m, tip radius 20 nm; Bruker, Mannheim, Germany) via a heterobifunctional benzaldehyde polyethylene glycol (PEG) linker (Broadpharm, San Diego, USA). Tip calibration was performed using the contact-based method within the SPM software. Sensitivity was determined by recording force-distance curves on a rigid surface submerged in an aqueous environment. Linear fitting of the repulsive region allowed conversion of voltage signals to displacement units. The spring constant of the AFM tip was calculated from thermal noise measurements. Topographical imaging of MEK cells was conducted in Quantitative Imaging (QI) mode with the following settings: setpoint = 0.5 nN, Z-length = 1.5 µm, and scan speed = 50 µm/s. Adhesion measurements were performed using force mapping mode controlled by SPM Control v.4 software (JPK Instruments, Berlin, Germany) with parameters: relative setpoint = 0.5 nN, Z-length = 1.5 µm, extend/retract speed = 10 µm/s, and contact time = 0.1 s. Regions of interest were selected along cell-cell contact, surface and leading edge. The scanned areas for force spectroscopy measurements covers 5 x 5 µm (25 x 25 pixels). Force-distance curve analysis was performed using JPK data processing software, providing metrics on interaction probability, binding strength, unbinding position and Younǵs Modulus.

### STED/AFM

STED/AFM experiments were done as described previously [14]. Briefly, the original STED microscope stage was replaced with an isolated Life Science stage equipped with a sample holder designed for inverted optical microscopes (Bruker Nano, Berlin, Germany). The NanoWizard 4 AFM scan head was mounted above this setup. Tip calibration was done as described in the AFM section. Next, the stained and living keratinocytes were positioned on the stage. Precise alignment of the AFM tip over the region of interest was achieved using the standard DirectOverlay™ protocol in the SPM software (JPK Instruments). DirectOverlay™ optically calibrates the AFM scan field with the optical microscope image by moving the AFM tip to nine predefined positions. At each of these positions, the optical microscope acquires an image (scan area: 30 × 30 µm; resolution: 1024 × 1024 pixels). The AFM software then analyzes these images to identify the tip’s position, enabling correction for optical distortions and image artifacts automatically. After alignment and calibration, STED images were acquired, followed by AFM topography imaging. The resulting datasets were superimposed within the SPM software, and adhesion force measurements were subsequently performed.

### Purification of recombinant E-Cad-Fc construct

Recombinant human E-Cad-Fc protein, comprising the full extracellular domain of E-Cad, was expressed in Chinese hamster ovary (CHO) cells. Once the cells reached confluency, the supernatant was collected, and the recombinant proteins were purified using protein A agarose affinity chromatography (Life Technologies, Carlsbad, USA). Protein purity was assessed by Coomassie staining and Western blot analysis.

### Generation of pSNAPf-mE-Cad construct

Complementary DNA (cDNA) was generated from a total RNA, derived from wild type (WT) MEK cells, though reverse transcription. Full-length cDNA of mouse E-cad was obtained by PCR amplification using as a template MEK-derived cDNA with following primers:

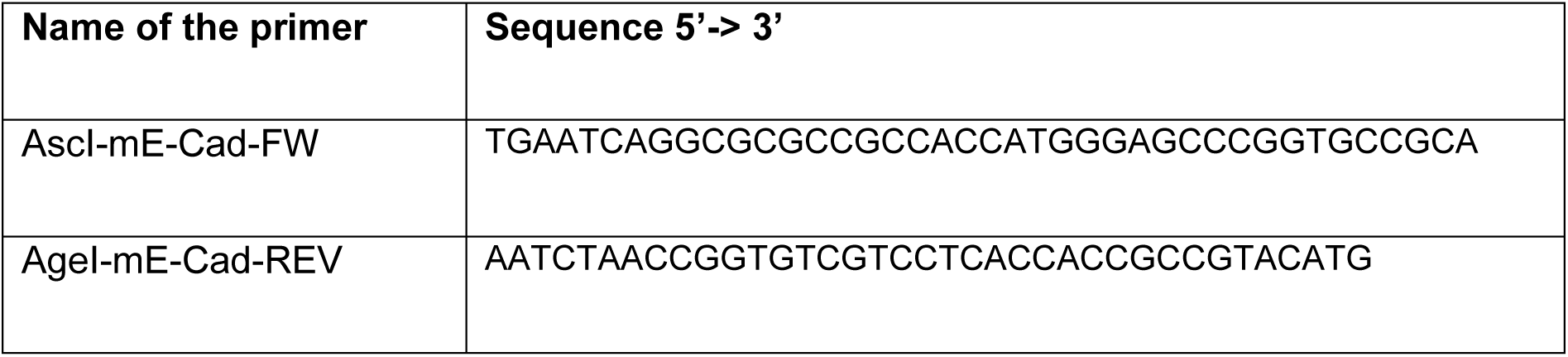

Within the forward primer (AscI-mE-Cad-FW), AscI restriction site was introduced. Similarly, the reverse primer (AgeI-mE-Cad-REV) carried in an AgeI restriction site. Both restrictions sites were used to insert the sticky-ended PCR product into linearized in advance pSNAPf vector (New England Biolabs, N9183S). The desired insert was fused upstream from SNAP-tag. Further restriction and sequencing analyses verified the construct.

### Data processing and statistics

For statistical significance between two groups a two-tailed Student’s T-test was applied. For comparison of more than two groups, analysis of variance (one-way ANOVA) followed by Bonferroni post hoc test, was performed. Error bars represent standard deviation. Results were considered as statistically significant when the p-values were equal or below 0.05. Figures were prepared with GraphPad Prism 8.

## Results

### E-Cadherin is present at the leading edge of migrating keratinocytes

To assess the expression of E-Cad during cell migration, we first performed scratch wound assays using (WT) murine keratinocyte and monitored wound closure over an 8-hour period. The scratch was done with a sterile pipette tip, and closure of the wound area was imaged at defined time points for up to 8 h (Figure 1 A). The quantification revealed that the scratch was reduced to approximately 60 % of its initial size after 6- and 8-hours post-scratch (Figure 1 B). Subsequently, we performed immunofluorescence staining for E-Cad and F-actin, as the latter was visualized with Alexa phalloidin. Images were taken at 0, 2, 4, 6, and 8 h post-scratch, with higher magnification views at the wound edge as well as in the following cells (Figure 1 C). We distinguished between the first and second row of cells next to the wound and quantified E-Cad fluorescence intensity. In the first row we observed a significant increase in E-Cad intensity 6 h after wounding, with a general upward trend detectable from earlier time points (Figure 1 D). In the second row, E-Cad intensity was moderately elevated from 2 h onward, although these changes did not reach statistical significance (Figure 1 D). To summarize, we found that E-Cad is present at the leading edge, representing the first cell row of migrating cells 6 h post-scratch.

**Figure 1:**
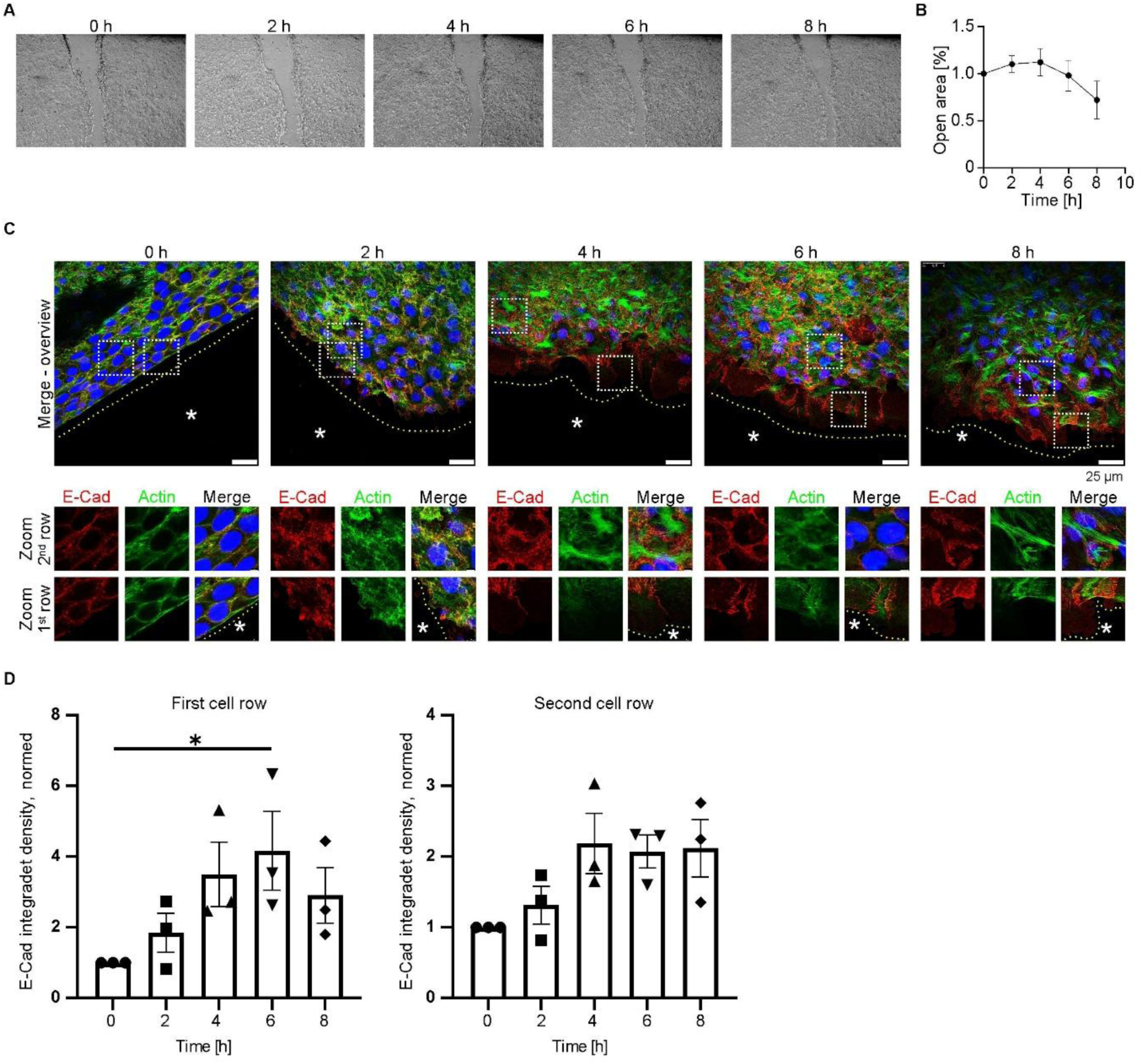
Scratch wound assay reveals elevated expression of E-Cad at the leading edge **A:** Murine keratinocytes were scratched with a pipette tip 24 h after the Ca^2+^ switch. Images were acquired at 0, 2, 4, 6, and 8 h post scratch. **B:** The cell-free area was measured over the 8 h period following scratch. **C:** E-Cad (red), F-Actin (green) and nuclei (blue) are visualized 0, 2, 4, 6, and 8 h after scratch at the wound edge. Zoomed-in images distinguish between the first (1^st^) and second (2^nd^) rows of keratinocytes adjacent to the wound. The leading edge of the firs cell row is indicate by a yellow doted line in the merge images. The scratched cell-free area is indicated by *. **D:** Quantification of E-Cad fluorescence intensity in the 1st and 2nd cell rows at 0, 2, 4, 6, and 8 h post scratch. N = 3, *p<0.05.

### E-Cadherin binding properties investigated with the STED/AFM-SMFS

Before investigating the E-Cad binding properties in migrating keratinocytes, we aim to measure the basal E-Cad binding properties in confluent keratinocytes. Therefore, we applied the combined STED/AFM-SMFS technique to probe specific E-Cad interactions at the cellular surface and at cell-cell contact sites. The principal STED/AFM setup is shown in previous publications [13, 14]. The AFM tip was functionalized with recombinant E-Cad protein. This technique allowed us directly to compare E-Cad binding properties in regions with overexpressed versus endogenous protein levels. For this purpose, we transiently transfected MEK cells with an E-Cad-SNAP construct. Prior to STED/AFM-SMFS experiments, cells were labeled with SNAP-CELL-TMR-STAR, enabling specific visualization of the tagged E-Cad protein. The E-Cad-SNAP constructs localized predominantly to cell-cell contact sites (Figure 2 A). High resolution STED imaging revealed a linear pattern of E-Cad staining at contact sites. Correspondingly, AFM topography imaging identified these cell-cell contact sites as elevated regions with raised incoming filaments. The STED signal for E-Cad aligned well with the topographical features observed with the AFM, confirming that the fluorescently labeled protein accumulates at the cell-cell contact (Figure 2 A). To investigate potential spatial variations, we performed SMFS measurements at cell-cell contact sites (cell borders) with those from regions above the nucleus (cell surface) (Figure 2 B). Analysis of binding frequency, binding strength and unbinding position revealed no significant differences between areas with overexpressed and non-overexpressed E-Cad, independent of the cellular area (Figure 2 B). Across all conditions, the binding frequency remained around 10%, indicating consistent interaction probability. The binding strength was slightly above 40 pN, and the unbinding position was slightly below 200 nm, independent of localization or expression level (Figure 2 B). These findings suggest that E-Cad maintains stable binding characteristics regardless of its local abundance or subcellular distribution.

**Figure 2:**
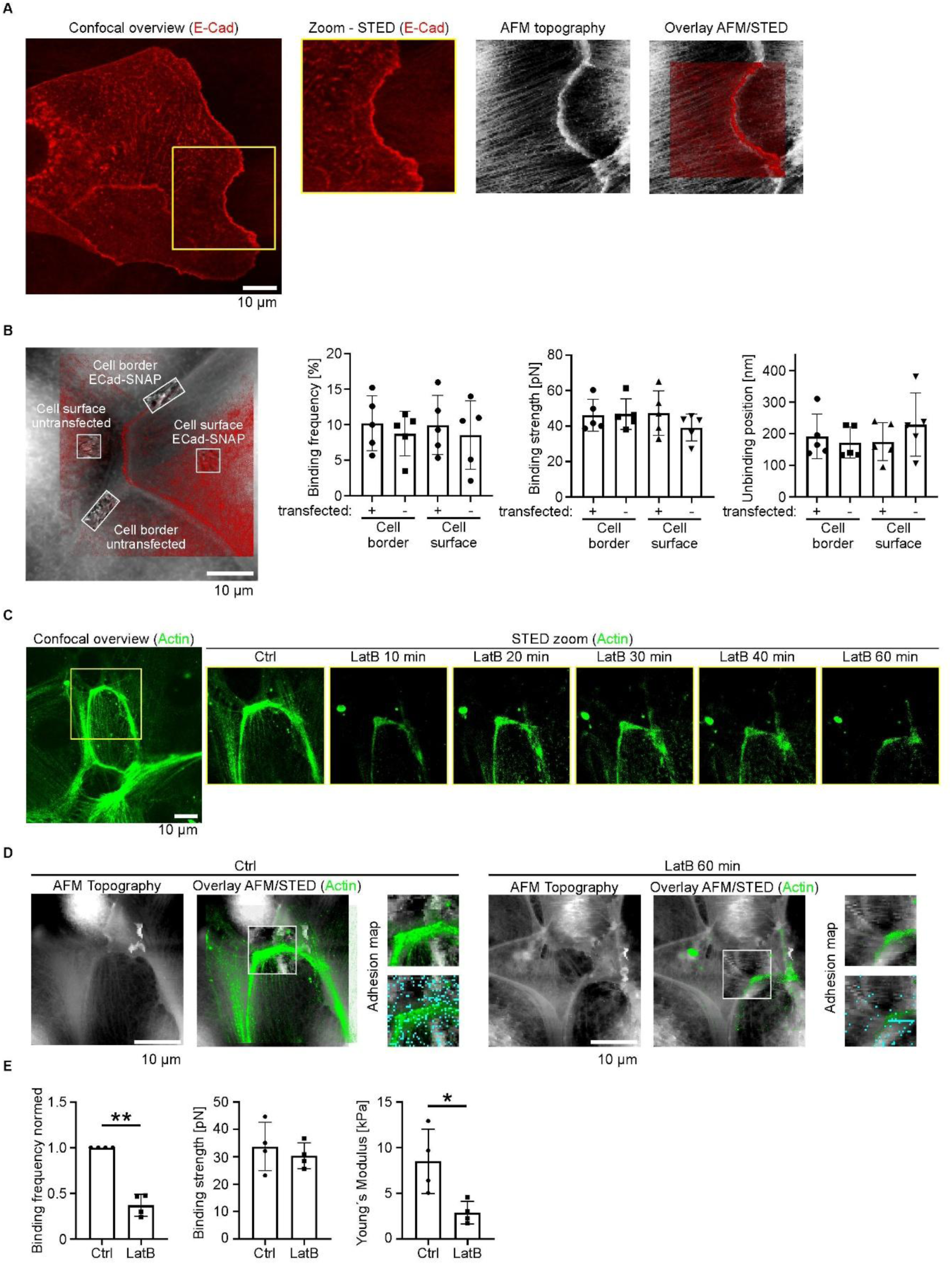
Investigation of E-Cad interactions by STED/AFM **A:** E-Cad-SNAP construct was overexpressed in murine keratinocytes and labeled with SNAP-CELL-TMR-STAR dye. Overlay images demonstrate that the topographical mapping overlaps the E-cad-SNAP signal. **B:** Binding probabilities were measured between an E-Cad-coated AFM tip and distinct cellular regions, including cell borders and cell surfaces, comparing cells with and without E-Cad overexpression. Quantification was done with 5 individual STED/AFM experiments, each consisting of two cell borders and surface area for transfected and non-ECad-transfected areas. **C:** Confocal imaging of F-actin labeled with Sir-actin following treatment with LatB for up to 1 h. **D:** STED/AFM imaging of murine keratinocytes labeled with Sir-actin. Cyan dots indicate E-cad binding events. **E:** Quantitative analysis of E-Cad binding events comparing untreated control cells to cells treated with LatB for 1 h; N=3; *p<0.05.

### E-Cadherin binding properties depend on the Actin cytoskeleton

To assess whether E-Cad binding is influenced by the actin cytoskeleton, we disrupted actin polymerization by using LatB, an inhibitor of actin polymerization. Cells were treated with LatB for 60 min. STED imaging revealed a continuous reduction of actin signal intensity within 1 h of LatB treatment, confirming the effective disruption of the cytoskeleton (Figure 2 C). Despite this, AFM topography imaging demonstrated that overall cellular morphology remained intact in the MEK cells, although STED imaging showed a strong loss of filamentous and cortical actin (Figure 2 D). SMFS measurements of E-Cad after 1 h of LatB treatment revealed a drastic reduction in binding frequency, indicating E-Cad binding is strongly dependent on the integrity of the actin cytoskeleton. Notably, the binding strength of the remaining E-Cad interactions was unaltered. Furthermore, LatB treatment led to a significant decrease in cellular stiffness, as indicated by a reduction in the Younǵs modulus (Figure 2 E). This supports the conclusion that actin polymerization is crucial not only for E-Cad-mediated adhesion but also for maintaining overall mechanical integrity of the cell.

### E-Cad binding properties at the leading edge of migrating keratinocytes

To directly assess the binding properties of E-Cad in migrating keratinocytes, we employed the STED/AFM approach. To create a defined wound area for migration, a culture-insert 2 well dish was positioned in the center of the AFM coverslip during cell seeding. The placeholder was removed 6 h prior to the STED/AFM experiment, allowing sufficient time for cells to migrate into the 500 µm gap and express E-Cad at the leading edge (Figure 3 A). We imaged the actin cytoskeleton at the leading edge by applying SirActin staining and probed E-Cad interactions at the single-molecule level. For this, AFM tips were functionalized with recombinant E-Cad to enable specific binding interactions. Confocal imaging at the wound edge revealed the presence of actin fibers oriented towards the gap. Corresponding AFM measurements at the same location showed increased rigidity in cellular regions with underlying cytoskeletal filaments, including actin, which are orientated towards the adjacent cell free space. Hence the AFM measures on the coverslip in the cell free area, this region shows a drastic elevated stiffness compared to the cellular tissue. These stiff cellular structures correlated with elevated topographical features seen in the AFM topography and with actin-protrusions clearly visualized in the STED/AFM overlay (Figure 3 B). These findings show that migrating cells can be investigated by the STED/AFM technique.

**Figure 3:**
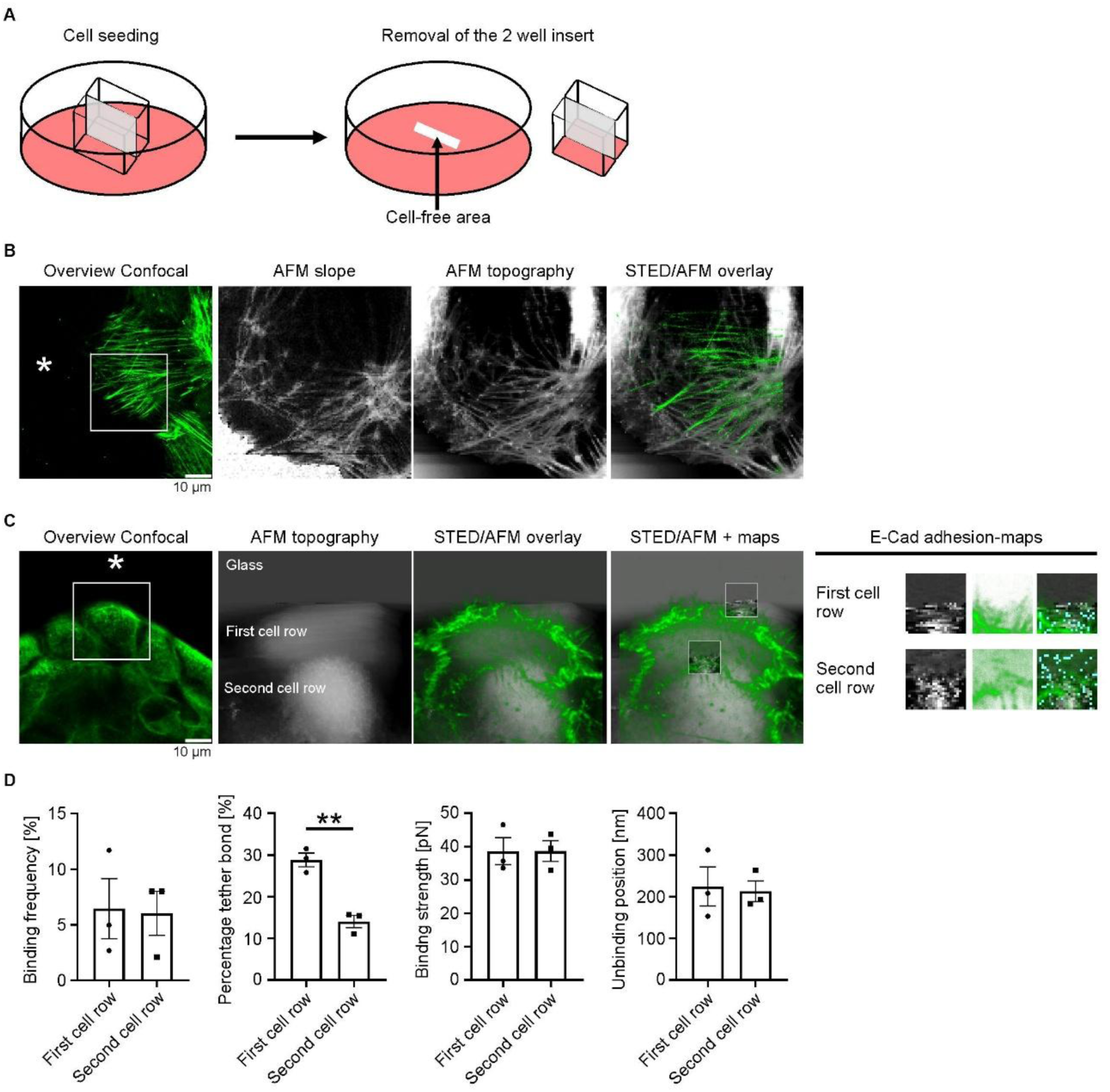
STED/AFM measurements on migrating keratinocytes **A:** Schematic illustration of the experimental setup. A two-well cell culture insert was placed on the coverslip within a 6-well plate. Removal of the insert led to 500 µm cell-free gap, allowing keratinocyte migration. **B:** STED/AFM imaging at the leading edge of migrating keratinocytes. Actin is labeled with Sir-actin. The cell-free area is indicated by *. **C:** STED/AFM imaging at the leading edge of MEK. E-Cad binding measurements were performed at both i.e the leading cell and at the first cell-cell contact. The AFM tip was functionalized with recombinant E-Cad protein. The scratched cell-free area is indicated by *. The measurements conducted at the cell-free area correspond to cover slip. **D:** Quantitative analysis of the E-Cad binding events where interactions at the leading edge were compared with those at the first cell-cell contact. N = 3, **p<0.01.

Finally, we performed E-Cad adhesion measurements using the STED/AFM technique at distinct regions of migrating keratinocytes: The first cell row and the second cell row, which already assembled cell-cell contacts (Figure 3 C). E-Cad binding events were detected already at the cellular surface of the first cell row, indicating that E-Cad proteins are present for interactions at the leading edge. Binding frequency was not increased in the second cell row directly behind. For calculation of the binding frequency, we subtracted the cell-free area. Interestingly, the proportion of tether bonds, which are /indication for cadherins weakly anchored to the cytoskeleton [14, 17], was significantly higher at the leading edge compared to the second cell row. In contrast, the binding strength and unbinding position did not alter between the two examined regions and remained consistent with the above presented measurements (Figure 3 D and see Figure 2 B), supporting the conclusion that E-Cad binding properties are conserved across migrating cell rows, despite the local differences in cytoskeletal anchorage.

## Discussion

In this study, we demonstrated that STED/AFM is a versatile technique for investigating keratinocyte migration, enabling both the measurement of cellular mechanical properties and single-molecule force spectroscopy of cadherins. We found that E-Cad is expressed at the leading edge of migrating keratinocytes and in this localization exhibits binding properties similar to those noticed in confluent cells. Using the combined STED/AFM approach, we further demonstrated that E-Cad binding depends on an intact actin network, which is itself essential for maintaining cellular stiffness. Furthermore, we showed that STED/AFM can be effectively applied to study the biophysical characteristics of migrating cells as well as the single molecule binding properties of their junctional molecules.

Scratch wound assays are widely used across various scientific fields, such as cell biology, cancer research, dermatology and regenerative medicine, to study cell migration, cell-cell interactions, proliferation and tissue regeneration [19–22]. The assay evaluates the rate of wound closure, which can be compared under different treatment conditions to assess the efficacy of therapeutic compounds. It is a relatively simple, cost-effective and easy-to-perform method for investigating wound healing and cellular migration. The technique mimics the *in vivo* process of cell migration during tissue repair. The main steps include creating a wound, typically by scratching a confluent cell monolayer with a pipette tip, capturing images at regular time intervals during the wound closure process, and analyzing these images to quantify the cell migration rate [20]. Scratch wound assays have also been applied in the desmosomal field, where it was shown that desmoglein 3 regulates the transition to a migratory keratinocyte phenotype through the inhibition of the p38MAPK pathway [23].

However, the mechanical properties and cadherin-binding properties of migrating keratinocytes have not been investigated before. Here, we demonstrate that the STED/AFM technique can be successfully applied to measure biophysical properties of migrating keratinocytes. We found that E-Cad expression was enhanced at cell-cell junctions at the leading edge 6 h after scratch. We investigated the binding properties with an E-Cad-functionalized AFM tip with overexpressed E-Cad proteins in MEK cells. The binding probability was approximately 10 %, consistent with cadherin interaction probabilities previously reported for living keratinocytes [16, 18, 24]. The measured binding strength was slightly above 40 pN, which is within the range described in earlier studies [25–27]. By applying the actin polymerization inhibitor Latrunculin B, we demonstrated that the frequency of E-Cad binding was drastically reduced, highlighting the crucial role of an intact actin network in supporting E-Cad interactions. In addition, we observed that disruption of the actin cytoskeletal network resulted in a loss of cellular stiffness, consistent with previous findings [28, 29]. This reduction in stiffness, reflected by a significant decrease in the Young’s modulus, indicates that actin filaments are key contributors to the mechanical integrity of cells[30].

Using the hybrid STED/AFM technique on migrating keratinocytes, cellular protrusions were visualized by live-cell STED imaging, which resemble more rigid and elevated structures observed in the AFM slope and height channel. Interactions between the E-Cad-coated tip and cellular binding partners revealed that molecules at the leading edge exhibit a higher percentage of tether bonds compared to those at the second cell row directly behind. This suggests that the binding partners at the leading edge are likely not firmly anchored and serve more as sensory, extra-junctional cadherins rather than primarily adhesive molecule [17, 31].

This study demonstrates the feasibility of combining STED/AFM with single molecule force spectroscopy measurements to investigate migrating cells, potentially bridging the gap between biophysical characterization and dynamic migration assays. This approach offers valuable insight into the mechanobiology of wound healing, an area that current methods can only partially address. Furthermore, the high-resolution STED imaging enables the correlation of mechanical measurements with cytoskeletal dynamics and protein localization within the migrating cells. In the context of cell-cell adhesion biology, this technique can be used to quantify adhesion forces at the molecular level and to investigate the dynamic remodeling of junctional complexes during migration. As demonstrated in this study, we measured the binding properties of E-Cad at the leading edge and compared them to those at stable cell-cell contacts. We found that both E-Cad pools exhibit similar binding properties, indicating that E-Cad molecules at the leading edge are functional and capable of mediating adhesion. For cancer research, this technique might offer a platform to study early mechanical changes associated with EMT and to identify biophysical properties of invasive cell behavior, for example downregulation of E-Cad. Thus, in this study we introduce a method to investigate the biophysical properties of single molecules during keratinocyte adhesion, which could lead to a better understanding of migration processes in tissue repair and wound healing.

## Author contributions

M.F. conducted experiments, acquired data, and analyzed data; M.F. and M.R. methodology,

M.F. designed the research study, M.F., M.R. and J.W. wrote the manuscript.

## Acknowledgments

We thank Martina Hitzenbichler for excellent technical assistance.

## Notes

### Competing Interest Statement

The authors have declared no competing interest.

